# An in-depth update on the benchmarks for 16S amplicon sequencing

**DOI:** 10.64898/2026.08.18.745548

**Authors:** Louis-Maël Guéguen, Alban Mathieu, Olivier Perin, Arnaud Droit

**Affiliations:** Département de médecine moléculaire, Faculté de médecine, Université Laval, Boulevard Laurier, G1V 4G2, Qúebec, Canada; Axe Endo-Nephro, Centre de recherche du CHU de Québec-Université Laval, Boulevard Laurier, G1V 4G2, Qúebec, Canada; Maasai team-Université Côte d’Azur, Inria, 2004, route des Lucioles, 06902, Sophia Antipolis, France; Digital transformation and innnovation, L’Oréal, 1 Avenue Eugène Schueller, Aulnay-sous-Bois, 93600, France

**Keywords:** benchmark, 16S amplicon sequencing, mock communities, metagenomics

## Abstract

Amplicon-based techniques provide a rapid and cost-effective approach for profiling microbial communities. However, the observed microbial diversity is influenced by a wide range of factors, encompassing pre-analytical steps such as the choice of primers and target regions, as well as the bioinformatic pipeline, including the selection of tools, reference databases, and parameter settings. Several benchmarks are already available in the literature, but the updates to important tools and databases, namely LotuS3, the Ribosomal Database Project and GreenGenes2, prompted our investigation. In this study, we conducted a comprehensive benchmark of the main bioinformatic tools and databases. Using seven regions for three publicly available mock communities of increasing complexity, we tested 38 possible combinations of sequence resolution algorithms (DADA2 stand-alone, LotuS3 (DADA2/UPARSE)), taxonomic classifiers and search tools (Kraken2, DECIPHER, RDP, MMseqs2, Lambda, and Metaxa2), and databases (SILVA, GreenGenes2, RDP, RefSeq, and Metaxa2). The region V1-V3, coupled with DADA2+MMseqs2+SILVA, DADA2+Metaxa2, or LotuS3 (DADA2)+RDP yielded the highest-quality estimates of the true diversity according to the metrics. We also demonstrated that even certain dominant genera remain difficult to detect, and that the quantification of all genera can be substantially over- or under-estimated, even when using optimal combinations of tools and reference databases.

## 1 Introduction

The 16S rRNA gene, which encodes the small ribosomal subunit, is ubiquitous among bacteria and archaea [1]. The balance between conservation and mutation rate that characterizes the nine hyper-variable regions (V1-V9) makes them suitable for classifying bacteria down to the genus level [2, 3], though resolution varies across regions and taxa. Using broad-range (near-universal) polymerase chain reaction (PCR) amplicon primers, researchers can target and amplify specific regions across most bacteria within a microbial community. The pool of amplicon reads obtained can be analysed to provide a comprehensive profile of the microbial community. This profile can be summarised by two primary outcomes: microbial identification and quantification.

The 16S analysis pipeline can be divided into several key steps: primer removal, read merging, sequence resolution into open or closed Operational Taxonomic Units (OTUs), chimera detection and removal, and taxonomic assignment. More recently, amplicon sequence variant (ASV)-based methods such as DADA2 [4] have enabled the inference of exact sequence variants. Unlike OTUs, ASVs are not the product of clustering, thus offering a finer taxonomic resolution. Both OTUs and ASVs can be labelled with a taxonomy through an assignment tool and a reference database.

There are numerous tools used for taxonomic assignment, based on different techniques. Alignment-based tools, such as Lambda [5] or BLAST [6], search for sequence similarity against a reference database via an index. In contrast, classifiers, such as Kraken2 [7] or RDP [8], rely on sequence properties (often *k* -mers) to assign taxonomic labels until the classification confidence falls below a specific threshold. Moreover, reference databases differ in their curation method, diversity, and scope. Among the widely used references are SILVA [9], GreenGenes [10], and RefSeq [11].

The variety of alternatives available at each step of the 16S analysis results in numerous possible combinations. Because updates to any component may impact the results, evaluating their performance against mock communities is essential to provide recommendations tailored to current tool and database versions for a given goal. Recently, three important projects were updated. The Ribosomal Database Project (RDP) [12] comprises both a taxonomic classifier and a reference, and received a major update reflecting 17 years of changes to its taxonomy and training set in 2024 [8]. GreenGenes was updated to its version GreenGenes2 (GG2) in 2024 [13] with a substantially expanded set of additional amplicon sequences, totalling 23 million sequences. LotuS2 was recently updated to version 3. Previous benchmarks focused on shotgun long-reads aligned to the 16S gene [14] or the sensitivity and specificity of ASVs and OTUs [15]. Since then, tools and databases have been updated. Thus, a new broader benchmark was necessary to provide the literature with updated recommendations identifying the top-performing pipeline.

In this study, we conducted a comprehensive investigation of multiple combinations across the 16S amplicon analysis workflow, from the 16S gene region to the taxonomic assignment, using three mock communities comprising 29 genera. The microbial identification and quantification were evaluated for each pipeline using standard metrics. By presenting results that extend beyond aggregated metrics, the benchmark should help researchers select among three pipelines depending on the objective and context of the study.

## 2 Materials and Methods

The materials and methods are summarised in Fig. 1. Figures were generated using R [16] and the R packages dplyr [17], tidyr [18], ggplot2 [19], cowplot [20], ComplexHeatmap [21], and ggh4x [22]. The Sankey diagram was generated using SankeyMATIC.

**Fig. 1.**
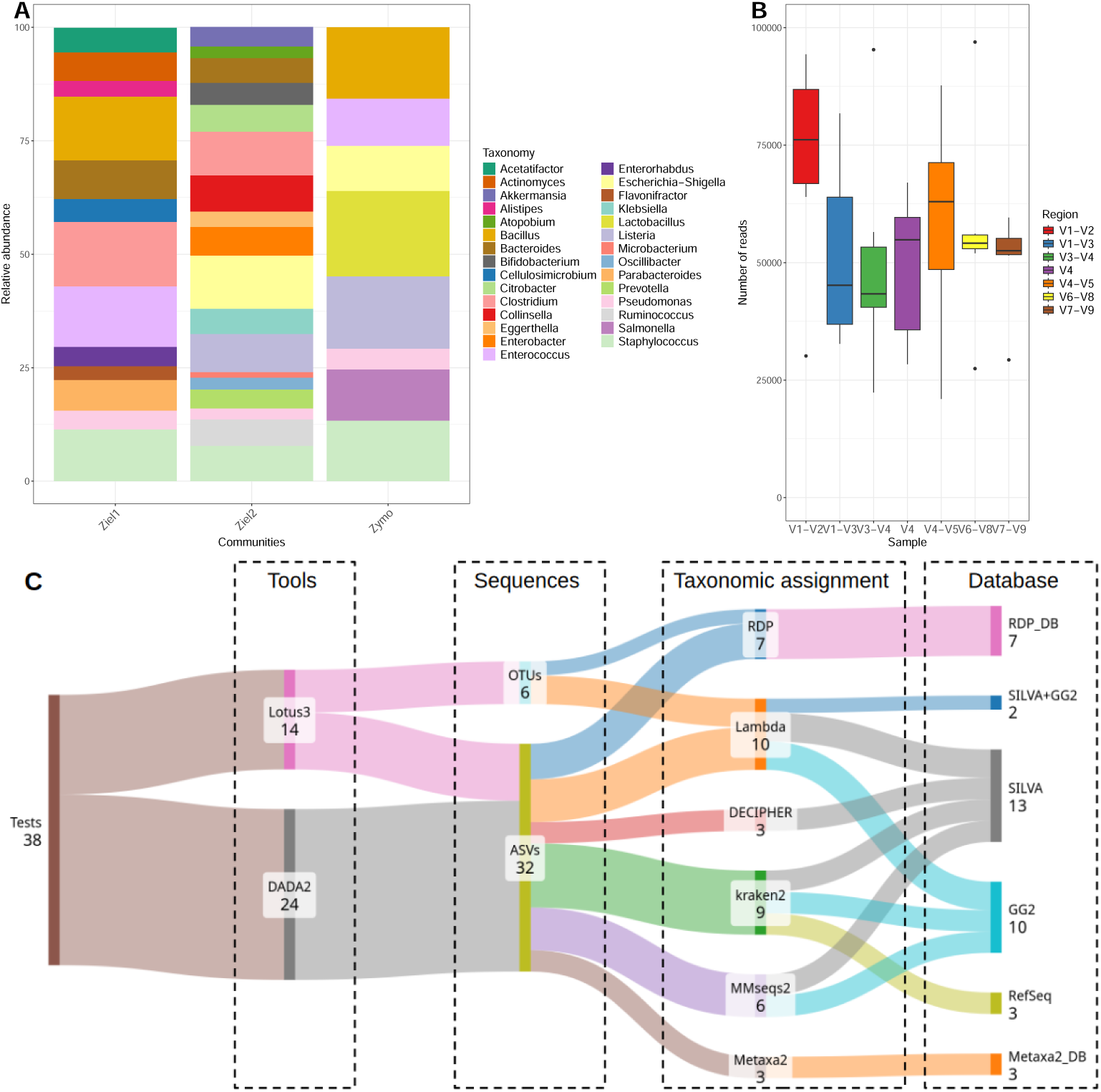
Overview of the benchmark settings. **A**. Genus composition of the three mock communities used in this benchmark (relative abundance in percentage). **B**. Sequencing depth for each region (scale limited to 100,000 reads, 2 data points not shown). **C**. Sankey diagram illustrating the tested combinations of tools and databases across all regions and communities. Each value represents the number of combinations using or yielding that specific component.

### 2.1 Mock communities

The theoretical compositions of the three mock communities from [23] are illustrated in Fig. 1, panel **A**. The Zymo community was composed of eight genera, ZIEL1 of 13 genera, and ZIEL2 of 18 genera. The communities were sequenced using the Illumina MiSeq technology, with paired-end reads of 300 bp, and a sequencing depth of approximately 50,000 reads (see panel **B**). The genera *Escherichia* and *Shigella* were grouped into *Escherichia-Shigella* based on the work of Lan and Reeves [24].

### 2.2 Tested combinations

The combinations were built to include the main sequence resolution algorithms, taxonomic assignment tools, and reference databases (Fig. 1**C**). These combinations can be divided into two groups: LotuS3 (v3.10)-based [25] and DADA2 (bioconda v1.34)-based. LotuS3 is designed to improve the ASV resolution by using only high-quality reads, and maps all reads for quantification. It provides an extensive range of options and tools, including for the resolution of ASVs (DADA2) or open OTUs (UPARSE), and the choice of taxonomic classifiers and databases. DADA2 is the most widely used tool for resolving ASVs.

LotuS3-based combinations used sdm as a primer removal tool (included in LotuS3), UPARSE [26] (as default setting) or DADA2 for sequence resolution, UCHIME3 [27] and LULU [28] for chimera detection, RDP (v2.14) or Lambda for taxonomic assignment. The databases used were SILVA (v138.1) and GreenGenes 2022.

DADA2-based combinations used Cutadapt v.5.2 [29] to remove the primers. The resolution of ASVs was tested using the three sequence resolution algorithms (”consensus”, ”pooled”, ”per-sample”). The taxonomy was assigned using one of the following tools: Kraken2 v2.12, DECIPHER v3 [30], MMseqs2 v18 [31], RDP v2.14, or Metaxa2 v2.2.3 [32]. The database for MMseqs2 and Kraken2 were built using the default parameters. The databases used were RDP v2.14, GG2 2024, RefSeq 2025 [33] and SILVA v138 for MMseqs2, and v138.2 for DECIPHER.

Tools that generate zOTUs or closed OTUs were excluded from consideration, as the field has shifted towards the use of ASVs. [34].

The combinations were tested on each hyper-variable region of each mock community. To evaluate the top-performing pipelines on a more complex dataset with a known ground truth, samples of the region V1-V3 from the three communities were collectively analysed as the ZIEL1-ZIEL2-Zymo (3Z) mock.

### 2.3 Percentage of reads used

The percentage of reads used by DADA2 was obtained by dividing the number of non-chimeric reads by the number of raw input reads (Eq. 1). For LotuS3, the number of reads in the matrix was divided by the number of raw reads (Eq. 2).

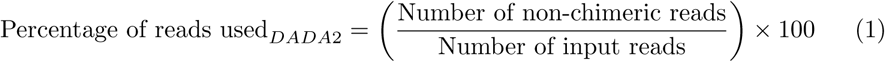

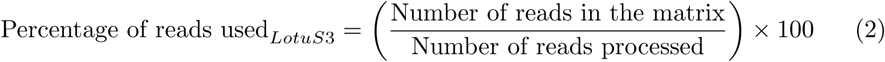

### 2.4 Metrics

The two key outcomes from each pipeline were the identification and quantification of genera. Two metrics derived from precision (Eq. 3) and recall (Eq. 4) were chosen to assess taxonomic assignment quality: the F1 score (Eq. 5) and the area under the precision-recall curve (AUPRC) (Eq. 6). Precision and recall are calculated using counts of true positives (TP), false positives (FP), and false negatives (FN). TP were genera from the mock community’s theoretical composition correctly identified in the predictions; FP were predicted genera absent from the theoretical composition; FN were genera from the theoretical composition of the mock community that were not present in the predictions. The F1 score was also illustrated as a function of the cumulative sum of abundance of the predicted taxa sorted by increasing abundance.

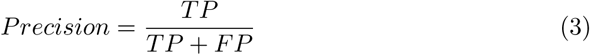

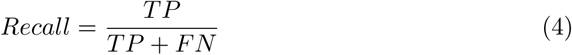

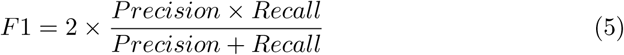

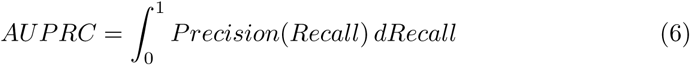

The Euclidean distance (L2, Eq. 7) was selected to assess the quality of microbial quantification. The theoretical proportions were multiplied by 50,000 (the average sequencing depth of the samples) to simulate theoretical counts. To allow comparison between theoretical and observed count matrices, genera missing in either the reference or experimental matrix were added with a pseudo-count of 1. The centred log-ratio (CLR) transformation (from the R package compositions [35]) was applied to both count matrices. The Euclidean distance (Eq. 7) was computed from the two transformed matrices. The transformation of theoretical relative abundances into pseudo-counts was chosen because it resulted in better contrast among pipelines by giving more weight to false positives and false negatives.

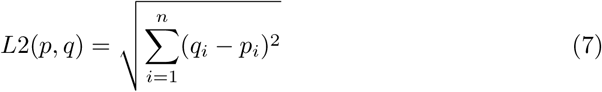

The metrics were computed using custom functions and the R package PerfMeas v1.2.5 [36].

### 2.5 Genus identification and under- or over-representation

To determine the identification frequency of microbes within each community, genus presence/absence in the taxonomic predictions was encoded as binary values (present = 1, absent = 0). The identification frequencies per community and per pipeline were computed using Eq. (8) and Eq. (9) respectively.

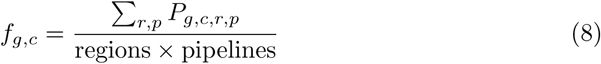

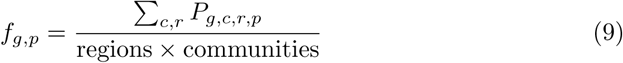

Where *P* was the binary-encoded *prediction*, *g* was *genus*, *r* was *region*, *c* was *community*, and *p* was *pipeline*.

To assess genus over- or under-quantification, the experimental raw counts were normalised to relative abundances. Missing genera were accounted for via zero-filling (relative abundance = 0). Then, the deviation from the theoretical composition was calculated as a percentage (Eq. 10).

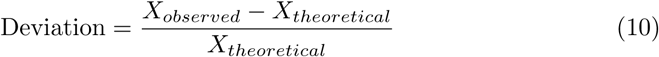

### 2.6 Runtime

The runtime was measured using wall-clock time from the bash command time. Both pipelines were given 30 threads to analyse the 3Z community on a machine equipped with dual AMD EPYC CPUs providing 32 threads each.

## 3 Results

### 3.1 Read usage and number of sequence variants

The percentage of reads used is shown by sample for DADA2 and by dataset for LotuS3 (Fig. 2). Runs of LotuS3 for region V7-V9 with DADA2 resulted in the complete filtering of the reverse reads, which yielded no results, in spite of using identical parameters to those applied to other combinations for the same samples. Despite filtering out more reads, DADA2 produced more sequence variants than LotuS3. LotuS3+UPARSE in particular yielded the lowest number of sequence variants, and the region V4 did not stand out as much as for the other combinations. Overall, ASV-producing pipelines were more sensitive compared to their OTU-producing counterparts.

**Fig. 2.**
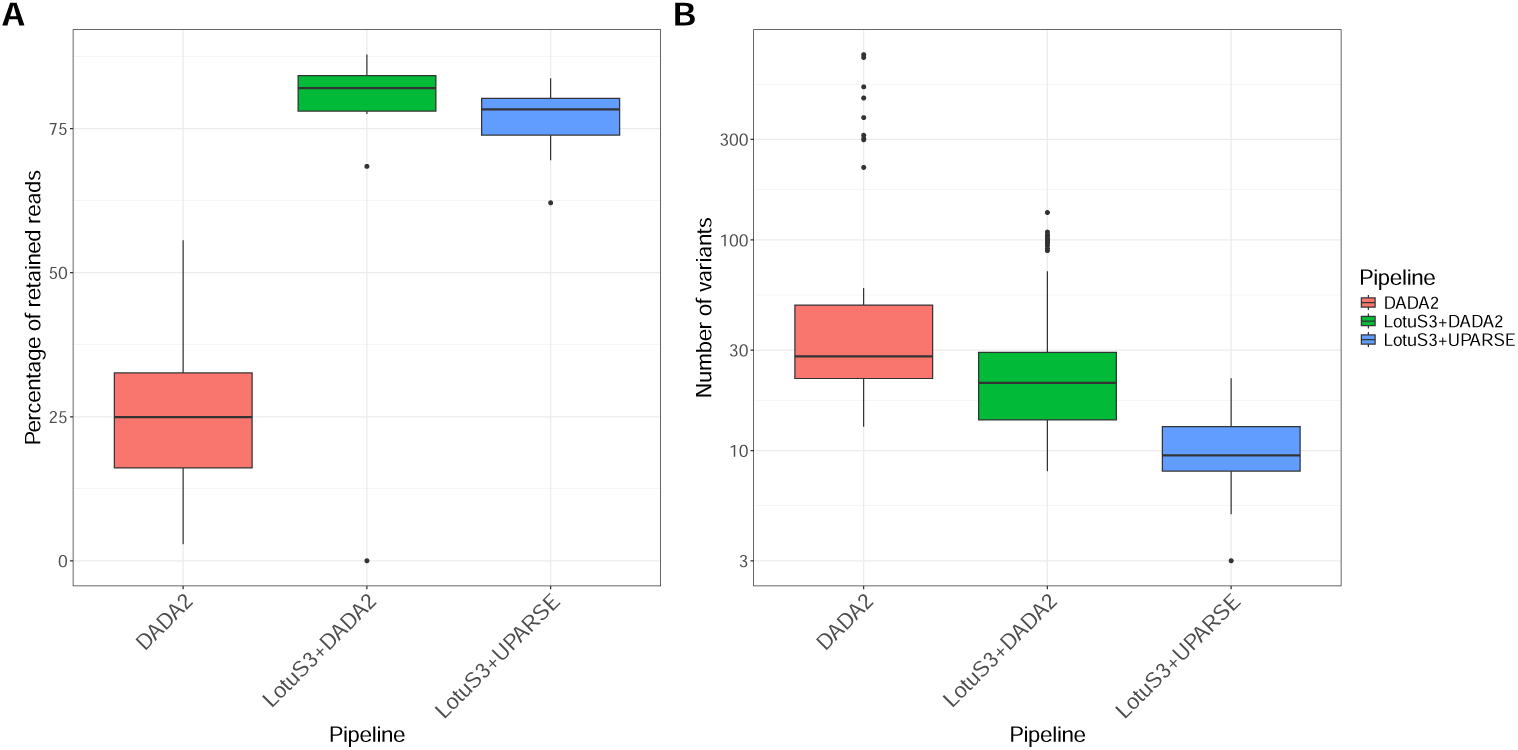
Read usage and variant production by pipeline. **A**. Percentage of input reads used. For DADA2, the values are the percentage of reads per sample. For LotuS3, the values are the percentage of reads per dataset. **B**. Number of sequence variants (ASVs, OTUs) produced by pipelines. The Y-axis is transformed with a log10.

### 3.2 The V1-V2, V1-V3, V3-V4 regions produced the highest-quality assessments

The quality of the results across all tested combinations was assessed by the three selected metrics (Fig. 3**A**). The metrics showed high variability for both pipelines. For LotuS3, the variability is not comparable for the region V7-V9, as results for LotuS3 (DADA2) were missing. For DADA2, the dispersion of the results observed (panels **B** and **C**) was due to the low precision and recall of the predictions produced by pipelines using DECIPHER or GG2. Additionally, a caveat of the L2 calculation method was discovered with DECIPHER. Pipelines yielding only ”uncultured” as a taxonomic assignment had a L2 distance of about 20, which was among the lowest results. The combinations including these tools were removed from downstream analysis. All results are illustrated nonetheless in Supp. Fig. 3. The three regions producing the highest-quality metrics were V1-V2, V1-V3, V3-V4 (Table 1), and were subsequently chosen for further analysis.

**Fig. 3.**
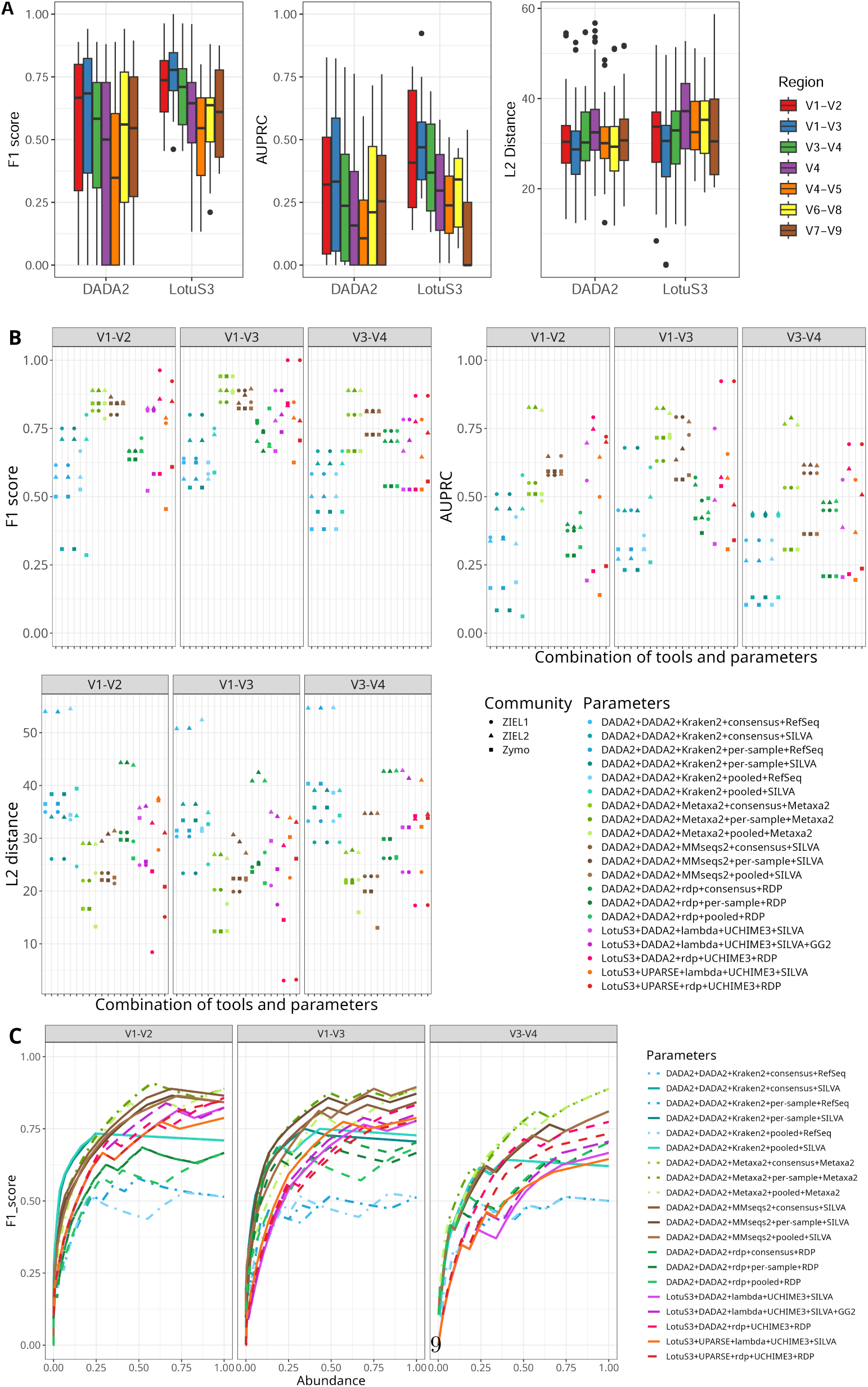
Metrics per region and pipeline. **A**. L2 distance, F1 score and AUPRC across regions of the 16S separated by family of pipelines. **B**. Metrics per pipeline for regions V1-V2, V1-V3 and V3-V4. **C**. F1 score as a function of the cumulative abundance of the mock community ZIEL2 (n=18).

**Table 1.** Metrics across regions. Values in bold are the best (highest or lowest) in their column. F1: median F1 score. L2: median of the Euclidean distance. AUPRC: median AUPRC.

| Region | F1 | L2 | AUPRC |
| --- | --- | --- | --- |
| V1-V2 | 0.667 | 31.1 | 0.345 |
| V1-V3 | **0.737** | **29.0** | **0.383** |
| V3-V4 | 0.645 | 31.6 | 0.265 |
| V4 | 0.56 | 34.5 | 0.193 |
| V4-V5 | 0.462 | 30.6 | 0.131 |
| V6-V8 | 0.625 | 29.8 | 0.256 |
| V7-V9 | 0.571 | 30.7 | 0.196 |

### 3.3 The four top-performing pipelines

Pipelines are detailed for the three best regions (Fig. 3**B**). The three methods of ASV resolution (”pooled”, ”consensus” and ”per-sample”) yielded similar metrics. Different *k* -mer values (*k* = 27, 21, 15) were tested during Kraken2 database construction for SILVA and GG2. However, the quality of the predictions dropped sharply (e.g. no predictions at the genus level) and thus the default values of the parameters were retained (k=35). Four pipelines had higher F1 scores, higher AUPRC and lower L2 distance than all other combinations: DADA2+MMseqs2+SILVA, DADA2+Metaxa2, LotuS3 (UPARSE)+RDP and LotuS3 (DADA2)+RDP (Fig. 3**B**). Finally, the highest

F1 score as a function of the abundance for the community ZIEL2 reached a maximum value of 0.89 for the combinations DADA2+Metaxa2 and DADA2+MMseqs2+SILVA coupled with the regions V1-V2 and V1-V3 (Fig. 3**C**). DADA2+Metaxa2 had the best performance for the region V3-V4 (F1 score = 0.89). The two LotuS3 pipelines had the best performance for the ZIEL1 community across the selected regions, with an F1 score reaching 1 for the region V1-V3 (Supp. Fig. 1**A**). The DADA2+Metaxa2 combination provided the highest-quality estimates (F1 score = 0.94) for the Zymo community across the selected regions (Supp. Fig. 1**B**).

### 3.4 Primer removal or retention

Primer retention improved results (L2 distance and F1 score) for the regions V3-V4 and V4, with LotuS3 (DADA2) (Supp. Fig. 4). While primer removal reduced the L2 distance for the regions V1-V2 and V1-V3, primer retention improved the F1 score for V1-V3. Primer retention had opposite effects on the alpha diversity (Shannon index) of pipelines producing ASVs and OTUs (Supp. Fig. 2). With UPARSE, the additional bases reinforces the clustering which reduces the number of variants produced, while they resulted in more variants with DADA2. This effect is stronger for regions V3-V4, V4, and V6-V8.

**Fig. 4.**
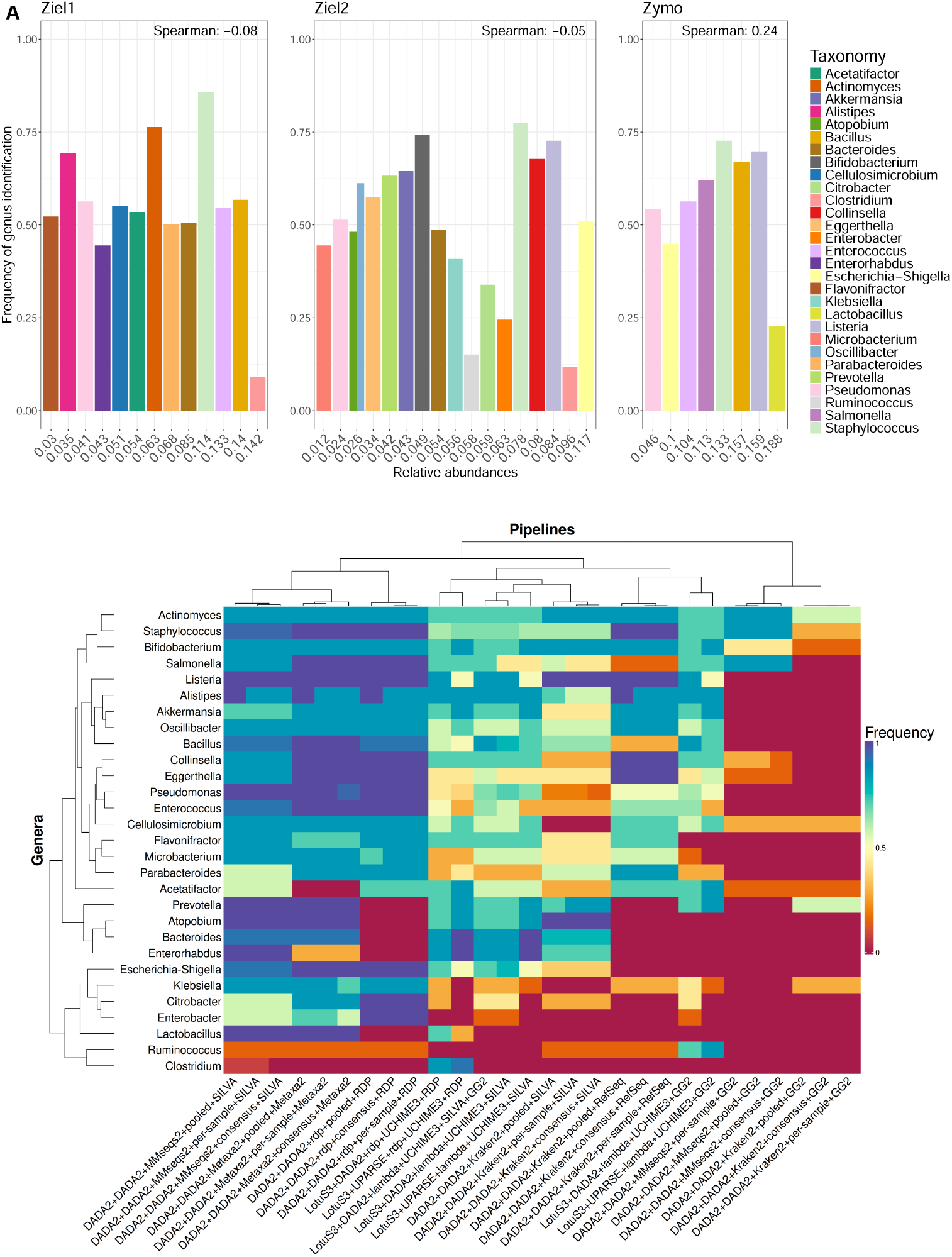
Identification of genera by community and pipeline. **A** Frequency of genera identification in the three mock communities. Genera are ordered by increasing theoretical abundance. The Spearman correlation between frequency and theoretical abundance of genera is shown for each community. **B** Frequency of genera identification per pipeline across regions and communities. The combinations using DECIPHER were removed for clarity. The hierarchical clustering trees of clusters and genera are based on Euclidean distance and complete method.

### 3.5 Examination of the most elusive genera across regions and classifiers

To refine the quality assessment, genera identification was investigated. The genera most frequently identified were *Staphylococcus* and *Actinomyces*, while the most elusive were *Clostridium* and *Ruminococcus* (Fig. 4**A**). Another genus that was difficult to detect was *Lactobacillus*, which was only identified using MMseqs2, Metaxa2, and LotuS3 (UPARSE/DADA2)+RDP (Fig. 4**B**). The correlations between the theoretical abundances and the frequency of identification were weak (Spearman, Fig. 4**A**).

### 3.6 Over- and under-quantification of genera across pipelines

The quantification of each genus was assessed for the four top-performing pipelines: DADA2+Metaxa2, DADA2+MMseqs2+SILVA, and LotuS3+(UPARSE/DADA2)+RDP in Fig. 5. The average across regions and communities is shown in panel **A**. The genera *Alistipes* and *Atopobium* were over-quantified by all pipelines. *Staphylococcus* and *Escherichia* were also over-quantified by pipelines with the exceptions of LotuS3+DADA2+RDP and DADA2+MMseqs2+SILVA respectively. The genera most accurately quantified by the DADA2 pipelines were *Bacillus*, *Bacteroides*, *Enterococcus*, *Oscillibacter*, and *Prevotella*. The pipelines based on LotuS3 accurately quantified *Bacteroides* and *Prevotella*. The quantification of genera is shown for regions V1-V2, V1-V3 and V3-V4 in panel **B**. *Atopobium* was the only genus that was strongly over-quantified in these regions. Overall, the microbial quantification largely diverged from the expected values across genera and pipelines.

**Fig. 5.**
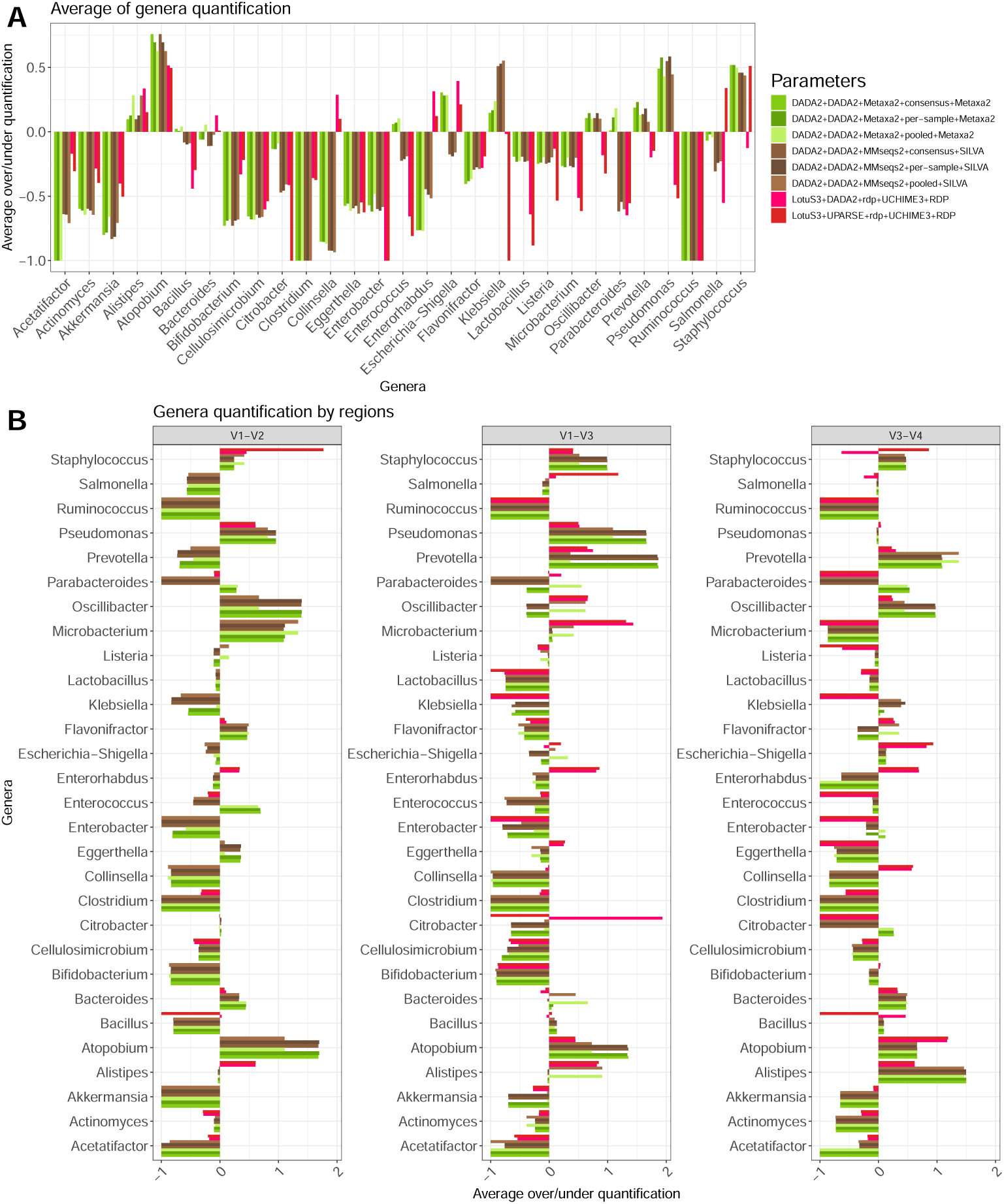
Deviation from the expected relative abundance of genera. **A** Average deviation from the expected relative abundance of genera across top-performing pipelines. A value of -1 indicates the absence of the genus, a value of 0 indicates the exact expected relative abundance, and a value of 1 indicates a two-fold relative abundance. **B** Deviation from the expected relative abundance of genera for regions V1-V2, V1-V3, V3-V4 across top-performing pipelines.

### 3.7 Further evaluation of the top-performing pipelines on the 3Z mock community

The 3Z community was analysed with DADA2+MMseqs2+SILVA and DADA2+Metaxa2 (both with ”pooled”, ”per-sample”, ”consensus”), and LotuS3+(UPARSE/DADA2)+RDP. The metrics are reported in Table 2. The ”pooled” algorithm provided the lowest L2 distance. While LotuS3 yielded the highest precision and F1 score, DADA2 (”pooled”) yielded the highest recall. Metaxa2 performed well and had the lowest distance but, unlike the other pipelines, did not identify the genus *Clostridium*. DADA2-based pipelines were substantially slower (9.5-fold) than the LotuS3-based ones (Table 2). Overall, each of the three ASV-producing pipelines yielded the best value for at least one metric.

**Table 2.**
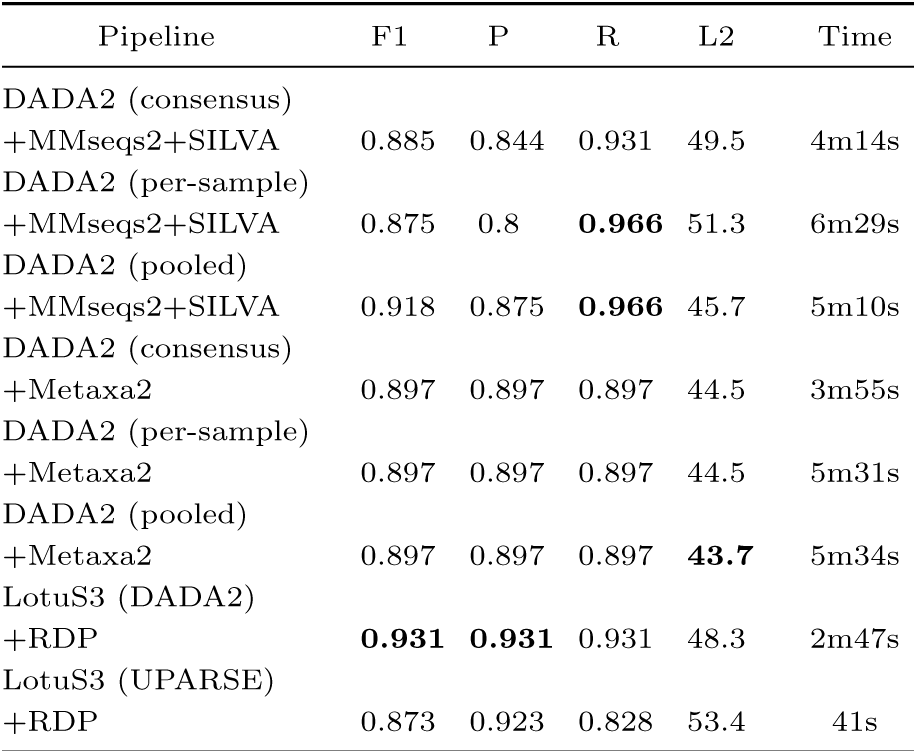
Metrics and runtime across pipelines for the 3Z mock community. Values in bold are the best (highest or lowest) in their column. F1: F1 score. P: Precision. R: Recall. L2: Euclidean distance. Time: wall-clock runtime.

## 4 Discussion

In this study, we comprehensively benchmarked 16S pipelines. The mock communities from [23] are a solid foundation as they contain a total of 29 different genera spanning a gradient of taxonomic complexities. The inclusion of seven regions increased the scope of the benchmark. Regions providing the highest-quality results were identified, then the top-four pipelines for these regions were tested on the 3Z community. Depending on the goal of the study, three pipelines are recommended.

### 4.1 Reasoning for the pipelines and metrics choices

The tools were chosen so that the benchmark was representative of the usage by the community. A few more were added to explore less widely used options such as Metaxa2 and MMseqs2. In the case of LotuS3, we had the choice to update certain tools and reference databases. RDP was updated to version 2.14 as it represented an important update of the taxonomy and was a drop-in replacement. Updating Green-Genes (2022) was also considered, but as the version in use was dated 2022 and its update may not reflect typical usage, we decided against it. Tools that generate zOTUs or closed OTUs were excluded from consideration, as the field has shifted towards the use of ASVs [34].

Performance metrics were selected as described by Simon *et al.* [14] to assess the quality of the microbial identification and quantification provided by the pipelines. Microbial identification was evaluated using two metrics that weight TP and FP, and FN differently, thereby refining the quality assessment. Results consisting only of ”uncultured” predictions were removed in Fig. 3 and beyond.

### 4.2 Read usage does not translate to more variants

There are several filtering steps in the 16S amplicon pipeline that aim to keep and use only the highest-quality reads for sequence resolution. DADA2 was more stringent than LotuS3 at this step, but the metrics did not reflect this lower usage rate of reads, which could be explained by (1) the low complexity of the mock communities (2) the higher number of variant produced. On a real microbiota, higher read usage may provide a more accurate quantification of genera, but the back-mapping step of LotuS3 relies on heuristics that may also skew the results.

### 4.3 Metrics

To narrow the comparison, we adopted a top-down approach: we first selected the regions that yielded the highest-quality results for the pipelines based on DADA2 and LotuS3. The performances of the two sets of pipelines showed similar patterns across regions. Following the selection of regions V1-V2, V1-V3, and V3-V4, individual performance of pipelines was evaluated. As the number of samples was limited, the similarity in quality of the DADA2 metrics across all three algorithms (”pooled”, ”per-sample”, or ”consensus”) was expected. The under-performance of GG2 in all combinations could be explained by the addition of millions of sequences specific to the V4 region. Such addition might have obscured sequence properties for *k* -mer-based methods and the indexes of alignment-based assignment tools. Then, the four top-performing combinations were assessed on the region V1-V3, which yielded the highest-quality results. The 3Z mock allowed a more challenging evaluation of the three algorithms for ASV resolution by using more samples of real data with a known composition. In addition to revealing more nuance in the results between chosen pipelines, this 3Z mock allowed us to suggest that ASVs yield better metrics than OTUs, consistent with previous work [15].

It was surprising to observe that primer retention could yield better results. The explanation might be that the additional bases help with the taxonomic assignment. Primer removal remains the recommendation as the gains were limited and specific to some regions.

### 4.4 Assessment beyond metrics

Beyond the global metric results, the benchmark tested the pipelines’ capacity to detect 29 genera. This allowed us to investigate whether some of the genera were more elusive than others and for which pipelines. The results illustrated in Fig. 4**A** suggest that sequence characteristics combined with tool heuristics may obscure abundant variants. Pipelines with good overall performance metrics could still miss dominant genera, some of which may represent important indicators in specific microbial environments. *Clostridium* is particularly relevant in studies of the gut microbiota, as species from this genus may cause botulism or tetanus. *Ruminococcus* is also an important genus of the gut microbiota. The absence of both or either of them from the outcomes of a human gut microbiome study would substantially skew the interpretation. Additionally, the microbial quantification across the 29 genera was overall far from the expected values. No pipeline was consistently more accurate than others across genera and the potential pathogenicity of the genera did not seem to be an indicator of over- or under-quantification. Such deviation could be attributed to primer bias, or copy number variation, as no control for that parameter was included in the pipelines. This aspect of the 16S amplicon analysis may still needs improvement. Together, the assessments of these two primary outcomes demonstrate the importance of evaluating pipelines beyond metrics to obtain a full understanding of the results.

### 4.5 Runtime

Apart from quality assessment, users with limited access to computational power may favour a given pipeline for its resource consumption. The bottleneck for CPU time was the sequence resolution, as the taxonomic assignment step was not constrained by the small size of reference databases and low number of variants. The design of LotuS3 allowed it to resolve sequence variants faster even when using DADA2 internally. RAM usage remained low for all pipelines.

### 4.6 Limits

This benchmark has several limitations. First, despite the analysis of the 3Z mock, the number of samples remained low. In particular, the method ”pooled” pools samples together prior to inference, while the two other algorithms perform inference per-sample. The complexity of the community also remained limited compared to that of naturally occurring microbial communities, and each genus was only represented by one species. Moreover, the three mock communities were sequenced using the same pairs of primers. Both facts limit the generalisation of this work. Second, we did not test all the combinations theoretically possible, as a component yielding low-quality results in several combinations is unlikely to perform substantially better in a different one. We also did not consider the merging step as a separate step in the benchmark as it is integrated into the pipelines. Finally, the merging of *Escherichia* and *Shigella* into a single genus favoured Metaxa2 over more recent work and made the resolution of our benchmark coarser. Taxonomic assignment is a difficult task, and the composition of the mock communities may have skewed the results in favour of certain taxonomic assignment tools and reference databases.

### 4.7 Recommendations

The benchmark tested 16S gene regions, sequence resolution algorithms, taxonomic assignment tools, and databases across three mock communities. The V1-V3 region provided the highest-quality estimates of microbial identification and quantification. Four pipelines outperformed the others for microbial identification: DADA2+MMseqs2+SILVA, DADA2+Metaxa2 and LotuS3 (UPARSE/- DADA2)+RDP. Given the higher performance of ASV-based methods and the age of the Metaxa2 reference (SILVA 111 [37] was released in 2012), we recommend either DADA2+MMseqs2+SILVA (”pooled” or ”per-sample”) or LotuS3+DADA2+RDP, both coupled with the primer pair used for V1-V3. In the specific case of a study aiming to accurately quantify genera that have not been impacted by the evolution of the taxonomy since 2012, DADA2+Metaxa2 is a suitable choice. Future research should focus on more accurate quantification of genera by systematically controlling for CNV.

## 5 Supplementary information

**Supp. Fig. 1.**
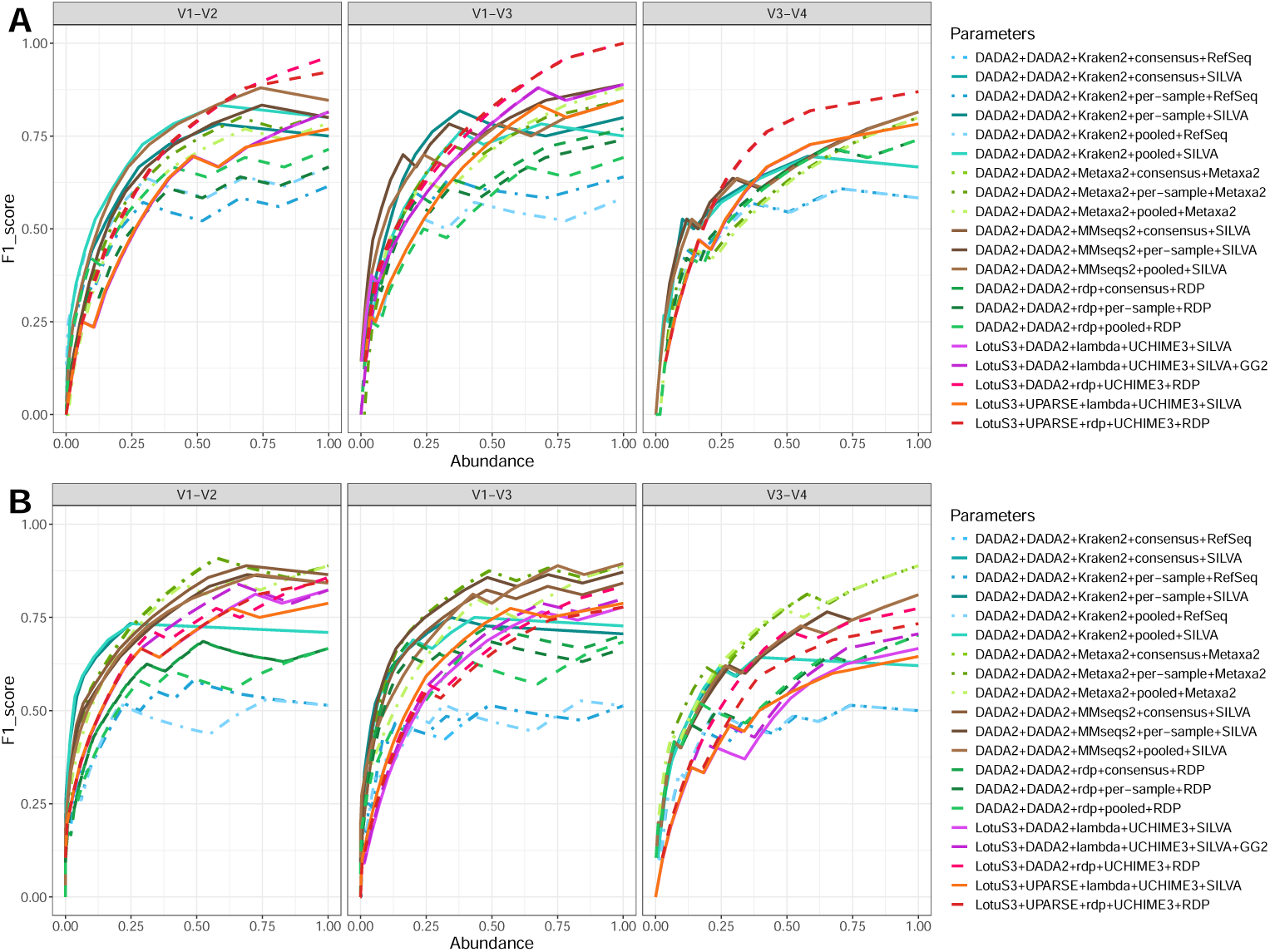
F1 score as a function of the cumulative relative abundance of genera. The x-axis is the cumulative relative abundance and the y-axis is the F1 score. **A** ZIEL1 community. **B** Zymo community.

**Supp. Fig. 2.**
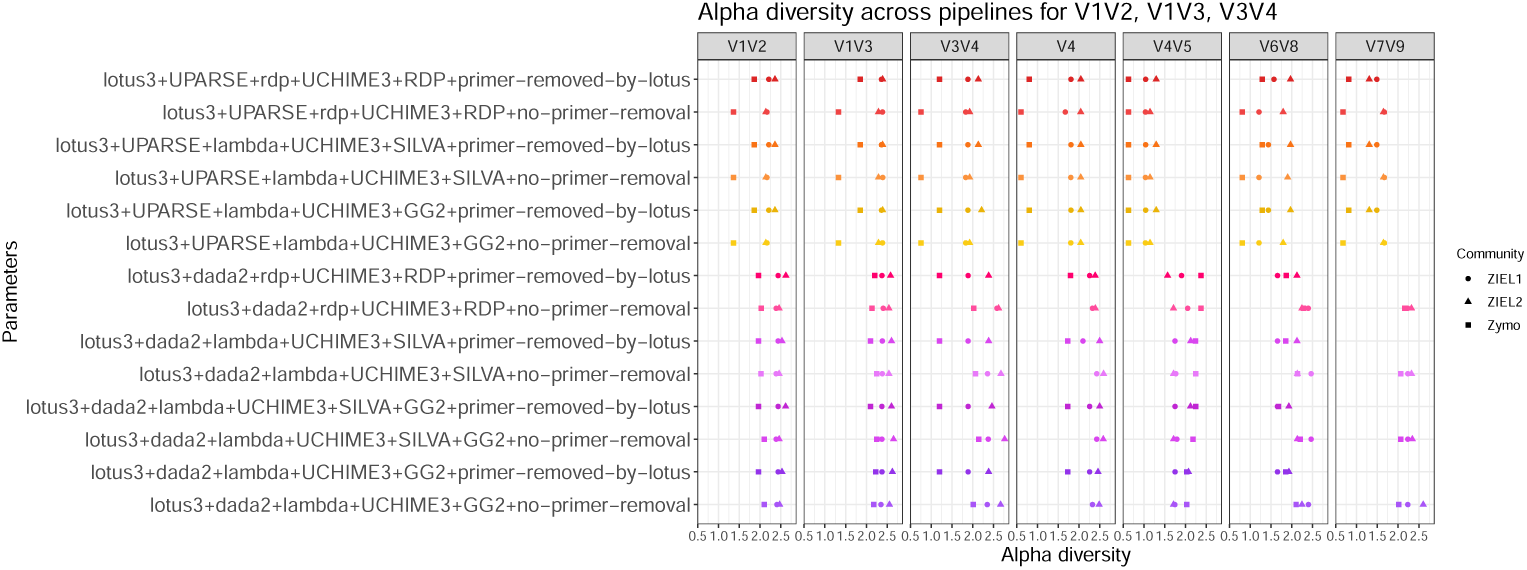
Alpha diversity (Shannon index) for each region for LotuS3 pipelines with and without primer removal by sdm (LotuS3).

**Supp. Fig. 3.**
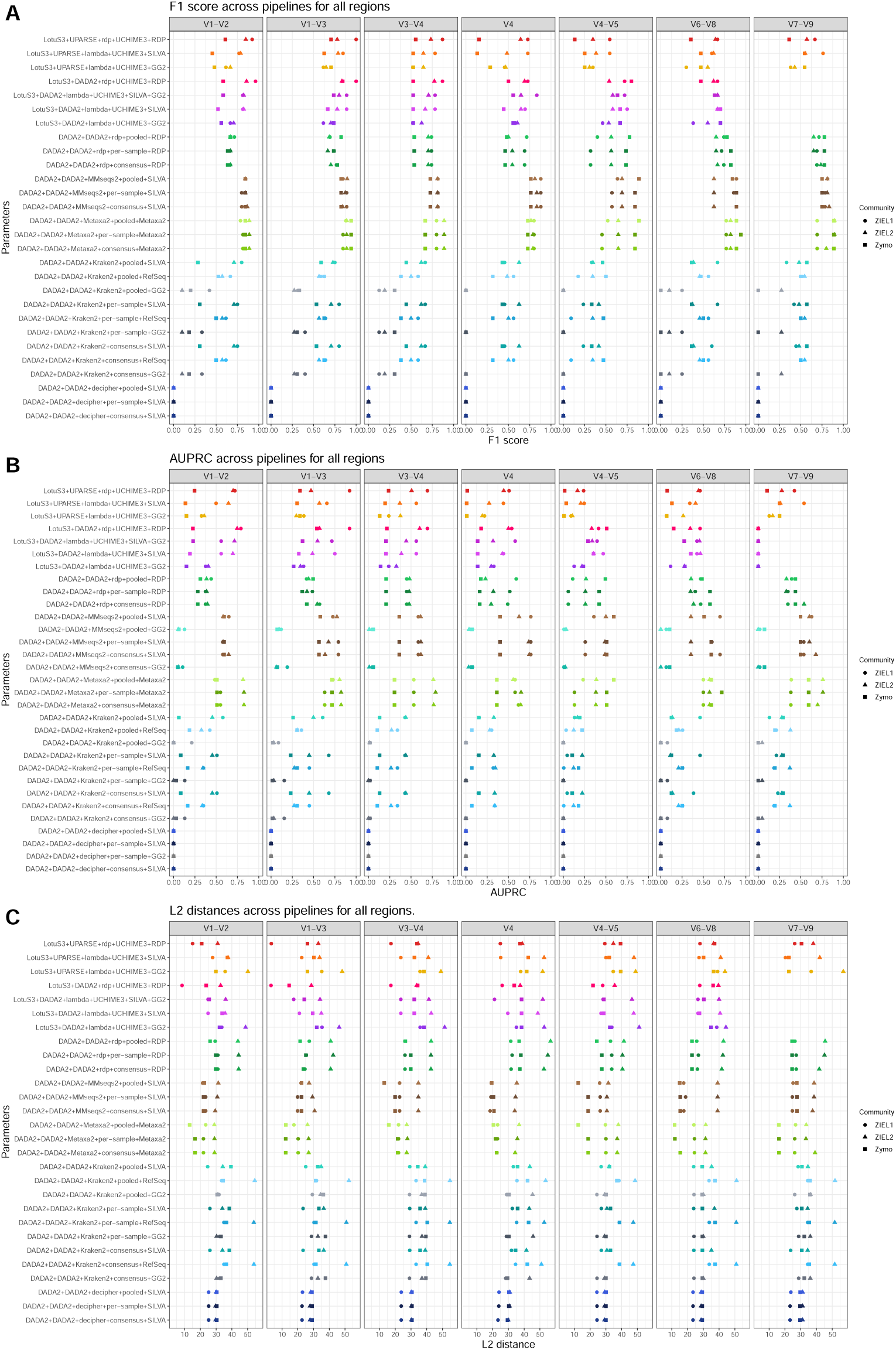
Metrics of all pipelines for all regions. **A** F1 score. **B** AUPRC. **C** L2 distance.

**Supp. Fig. 4.**
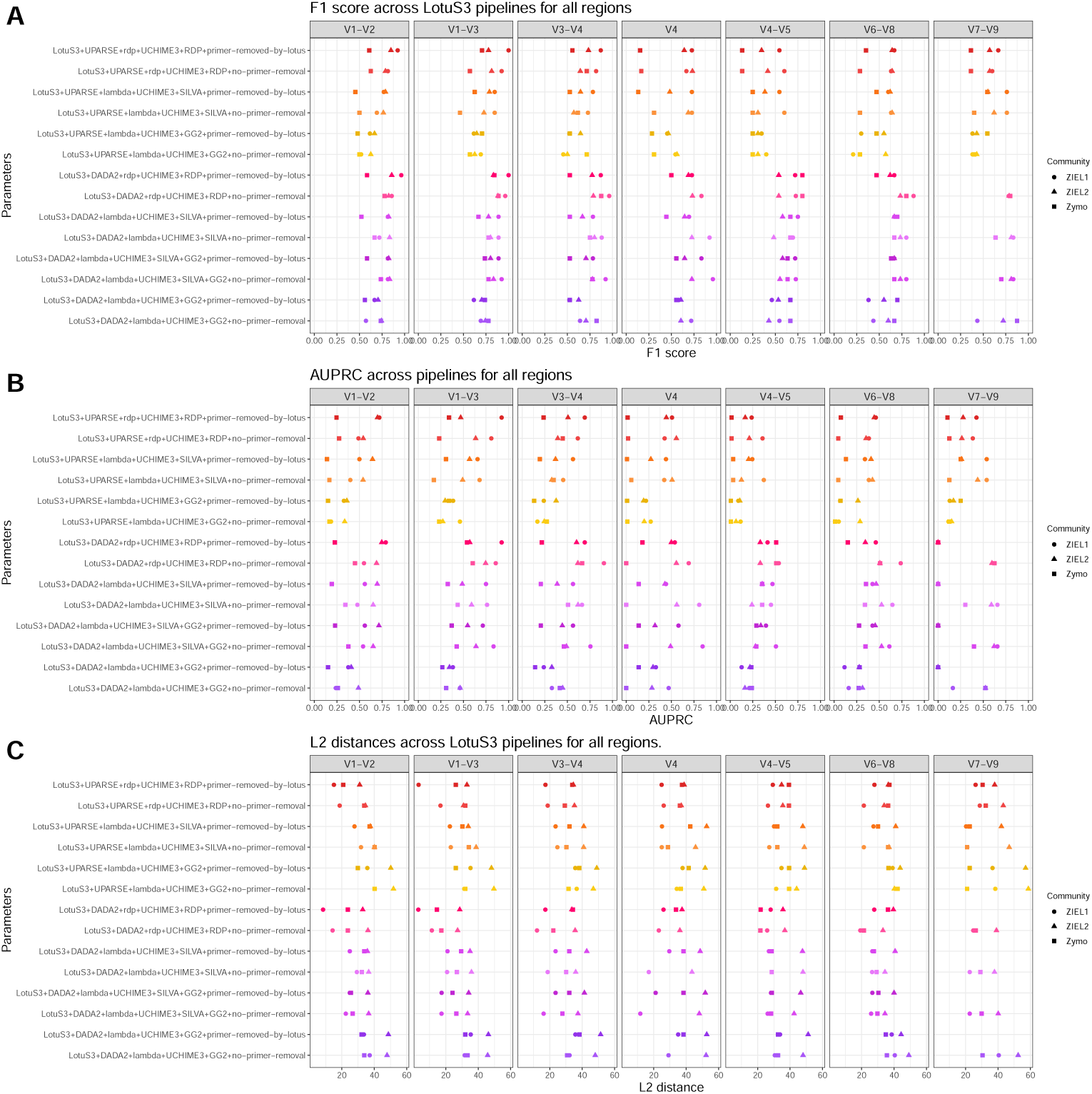
Comparison of the metrics between LotuS3 pipelines with and without primer removal. **A** F1 score. **B** AUPRC. **C** L2 distance.

## Acknowledgements

We thank Falk Hildebrand for helping with LotuS3. The first author would like to thank Audrey Lemaçon for her comments on the manuscript, and Romain Durand for his proofreading of the article.

## Declarations

### Ethics approval and consent to participate

Not applicable

### Consent for publication

Not applicable.

### Data availability

Data can be found at PRJNA674596.

### Code availability

Scripts used to run the benchmark, to generate figures and tables, as well as the figures are available on [GitHub](https://github.com/Louis-MG/16S_benchmark) and [Zenodo](https://zenodo.org/records/21267173).

### Author contribution

Conceptualization: Arnaud Droit; funding acquisition: Arnaud Droit; formal analysis, investigation: Louis-Maël Guéguen, methodology: Louis-Maël Guéguen, Alban Mathieu; writing – original draft; Louis-Maël Guéguen; writing – review & editing, all authors: Louis-Maël Guéguen, Alban Mathieu, Arnaud Droit.

